# Interspecific plastome recombination reflects ancient reticulate evolution in *Picea* (Pinaceae)

**DOI:** 10.1101/097519

**Authors:** Alexis R. Sullivan, Bastian Schiffthaler, Stacey Lee Thompson, Nathaniel R. Street, Xiao-Ru Wang

## Abstract

Plastid sequences are a cornerstone in plant systematic studies and key aspects of their evolution, such as uniparental inheritance and absent recombination, are often treated as axioms. While exceptions to these assumptions can profoundly influence evolutionary inference, detecting them can require extensive sampling, abundant sequence data, and detailed testing. Using advancements in high-throughput sequencing, we analyzed the whole plastomes of 65 accessions of *Picea,* a genus of ~35 coniferous forest tree species, to test for deviations from canonical plastome evolution. Using complementary hypothesis and data-driven tests, we found evidence for chimeric plastomes generated by interspecific hybridization and recombination in the clade comprising Norway spruce (*P. abies*) and ten other species. Support for interspecific recombination remained after controlling for sequence saturation, positive selection, and potential alignment artifacts. These results reconcile previous conflicting plastid-based phylogenies and strengthen the mounting evidence of reticulate evolution in *Picea.* Given the relatively high frequency of hybridization and biparental plastid inheritance in plants, we suggest interspecific plastome recombination may be more widespread than currently appreciated and could underlie reported cases of discordant plastid phylogenies.

## Introduction

Technical feasibility coupled with an apparently simple mode of evolution have made plastid genomes (plastomes) a workhorse of plant systematics. While nuclear genomes vary in size by orders of magnitude, a typical land plant plastome contains around 115 unique genes distributed over 120-160 kbp (Bock 2007). Plastid genes are generally conserved and encode the two photosystems, the large subunit of the carbon fixing enzyme RuBisCO, and ~30 other housekeeping proteins. Compared to the nuclear genome, plastome mutation rates are characteristically low and coding sequences may be subject to stronger functional constraints (Wolfe et al. 1987). In addition, the uniparental plastid inheritance found in most species is expected to result in clonal lineages, a smaller effective population size (*N_e_*) than the nuclear genome, and the absence of sexual recombination.

Exceptions to nearly every assumption of this simple model of plastome evolution are known (reviewed in Wolfe & Randle 2004). Of these, biparental plastid inheritance is particularly widespread phenomenon. Plastids are most often transmitted by the seed parent in angiosperms but about 20% of species have plastids present in mature pollen, which indicates the potential for biparental inheritance (Corriveau & Coleman 1988; Zhang et al. 2003). In coniferous gymnosperms, where plastid inheritance is typically paternal, realized biparental inheritance has been documented in *Pinus* (White 1990), *Picea* (Sutton et al. 1991), and *Larix* (Szmidt et al. 1987). Heteroplasmy resulting from biparental inheritance can be expected to be transient due to the vegetative sorting of organelles at every cell division (Wolfe & Randle 2004). However, a recent study of whole plastomes in *Citrus* revealed heteroplasmy consistent with hybridization in germplasm from mature trees (Carbonell-Caballero et al. 2015). Biparental plastid inheritance and stable heteroplasmy allow for violation of another axiom of plastome evolution: sexual recombination.

Robust evidence for recombination between biparentally inherited plastomes in natural populations is lacking. However, population studies of *Pinus* (Marshall et al. 2001) and *Cycas* (Huang et al. 2001) have invoked recombination to explain unusual patterns of plastome polymorphism and linkage disequilibirum. Interspecific recombination has been hypothesized as the source of the markedly different phylognies inferred from different plastid loci in *Silene* (Erixon & Oxelman 2008), *Picea* (Bouillé et al. 2011), and the diatom genus *Pseudo-nitzschia* (D’Alelio & Ruggiero 2015). Given that all sites in the plastome should share the same topology under clonal inheritance, recombination may more broadly underlie cases of discordance among plastid loci in studies where its presence has not been inferred (e.g. polygrammoid ferns, Schneider et al. 2004; Asteraceae, Doorduin et al. 2011). As recombination can severely bias population and phylogenetic analyses (e.g. Schierup & Hein 2000), new detection methods are warranted. In particular, approaches using complete chromosomes or organelles can exploit the expected spatial mosaic of evolutionary histories generated by recombination (e.g. Maynard Smith 1992) and use structural features to identify probable breakpoints or hotspots.

Repetitive sequences contribute to intra- and intermolecular recombination during plastome replication and repair (Day & Madesis 2007; Maréchal & Brisson 2010). During intermolecular recombination, repeats can act as recognition or cleavage sites through primary and secondary structures (Kawata et al. 1997). In addition, repeats may help maintain plastome size and content during recombinational DNA repair. For example, double-stranded breaks in *Chlamydomonas* plastomes are repaired through homologous recombination, which can result in an unstable plastome dimer that is reduced through intramolecular recombination between repeats (Odom et al. 2008). This process could explain why sexual recombination hotspots in the *Chlamydomonas* plastome are correlated with dispersed and inverted repeats (Lemieux et al. 1990; Odom et al. 2008). Based on this evidence, we reasoned rare sexual recombination in land plant plastomes should also be associated with repeats.

We developed and applied phylogenetic tests for recombination to 65 whole plastomes representing all ~35 extant species of *Picea,* a genus of coniferous trees broadly distributed in the northern hemisphere (Farjon 1991; Eckenwalder 2009). In contrast to the canonical quadripartite plastome structure found in most plants, the large inverted repeat (IR) region is highly reduced in *Picea* and other conifers (Strauss et al. 1988; Wu et al. 2011). Instead, the *Picea* plastome contains three single copy regions (the F1, F2, and F3 regions), which are intervened by three large repeats (Wu et al. 2011, see Fig. 1). Large structural rearrangements between these repeats have been reported (Wu et al. 2011; Nystedt et al 2013), which supports their role as recombination substrates during intra- and intermolecular recombination. We predicted sexual recombination in *Picea* would generate different evolutionary histories among the F1, F2, and F3 regions – regardless of genomic context, mutation rates, or natural selection – but not within them. Combining this hypothesis-based approach with purely data-driven tests, we identified at least one interspecific recombination event that produced a chimeric plastome inherited by over one-third of *Picea* species. This result remained after controlling for process that may generate patterns similar to recombination, such as process heterogeneity, saturation, and positive selection. Given that hybridization occurs in ~15% of plant genera (e.g. Ellstrand et al. 1996, Whitney et al. 2010), such recombinant plastomes may not be unique to *Picea.*

**Figure 1.**
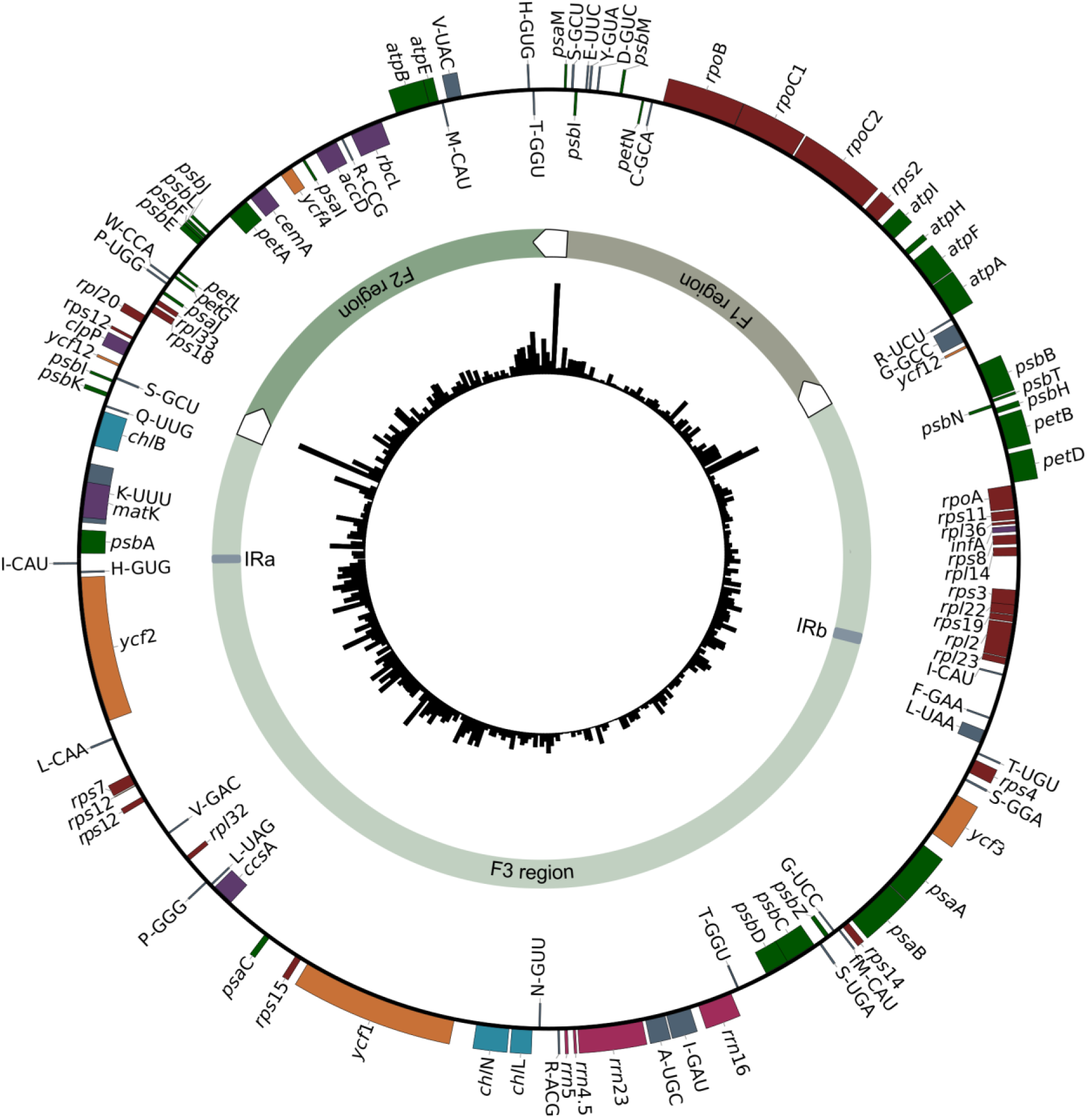
Consensus map of the *Picea* plastome. The outermost ring displays genes colored by functional group: genes on the inside and outside of the circle are transcribed in the clockwise and counterclockwise directions, respectively. The second circle displays the boundaries of the three major structural regions (F1, F2, and F3) of the plastome, which are intervened by large repeats represented by white arrows (not to scale). The innermost circle depicts average pairwise nucleotide diversity (π) estimated in 350 bp moving windows.

## Results

### Plastome assembly and divergence

We used whole-genome shotgun sequencing to generate 61 draft plastomes, which together with existing assemblies represent all 35 commonly-recognized *Picea* species. Total genomic DNA was sequenced on three different Illumina platforms at differing multiplexing levels and read lengths (See Materials and Methods), but all assemblies had high coverage and large scaffolds (Table 1). *De novo* plastomes averaged 121,885 bp (± 2,861) or about 98% of reported reference-quality assemblies (Nystedt et al. 2013; Jackman et al. 2015; Wu et al. 2011; Yang et al. 2016). Assemblies generally comprised five large non-overlapping scaffolds, which correspond to the intervening regions between the two copies of the highly-reduced inverted repeat (IRa, IRb) and the three large Pinaceae-specific dispersed repeats (Fig. 1). Plastomes assembled from two paired-end libraries were more contiguous despite lower sequencing depth and shorter reads (Table 1), possibly because of improved resolution of repetitive sequences (Prjibelski et al. 2014) or differences in Illumina chemistry. Assembly statistics, detailed locality information, and Genbank and Short Read Archive (SRA) accession numbers for each individual are reported in Supplementary Table S1.

**Table 1.** Sequencing and assembly metrics for 61 *Picea* draft plastomes. Standard errors are reported in parentheses.

| Sequencing system | Mean number of scaffolds | N50 scaffold length (bp) | Average largest scaffold (bp) | Mean read depth |
| --- | --- | --- | --- | --- |
| *Illumina HiSeq 2000 & 2500*<br>2x100 bp, 2x125 bp | 5.77 (1.76) | 56,475 (2,528) | 66,112 (13,562) | 46.83 (23.32) |
| *Illumina HiSeq 4000*<br>2x150 bp | 6.75 (1.35) | 22,939 (1,162) | 55,847 (6,949) | 413.51 (239.18) |
| *Illumina MiSeq*<br>2x100 bp | 7 | 23,241 | 55,799 | 47.17 |

Subsequent analyses included four additional published *Picea* plastomes (Nystedt et al. 2013; Jackman et al. 2015; Wu et al. 2011; Yang et al. 2016), yielding a total of 65 assembled genomes. Spruce plastomes were highly similar, averaging 97.9% sequence identity in pair-wise comparisons. Gene content was conserved and comprised 74 protein-coding genes, 36 tRNA genes, and four rRNAs. No evidence of rearrangements within scaffolds was found, but we were unable to evaluate structural changes spanning multiple scaffolds, such as the inversion of the F2 region between *P. morrisonicola* and *P. abies* (Nystedt et al. 2013). Indels were found in only three genes: *acc*D, *chl*N, and *ycf1.*

Nucleotide diversity (π) across the plastome was relatively low (π = 0.0030; Table 2). Sliding-scale estimates of π in non-overlapping 350 bp windows revealed three ‘hot-spots’ of diversity occurring near the boundaries of the structural regions (Fig. 1). Five genes *(psbH, psbT, rpl23, ycf1, ycf2)* were more polymorphic than intergenic spacers on average. Notably, *ycf1* and *ycf2* together contain 57% of all coding sequence polymorphisms (Supplementary Table S2). Mean synonymous and non-synonymous divergence was low (*d*N = 0.017 ± 0.024, *d*S = 0.040 ± 0.031) but varied markedly among genes (Fig. 2). Mean substitution rates were significantly elevated in the *ycf* genes (GLM, *t* = 2.828, p < 0.001) – a pattern driven entirely by *ycf1* and *ycf2* – and in the photosystem II complex *psb* (GLM, *t* = 2.103, p < 0.05; Fig. 2). Six genes comprised solely non-synonymous substitutions while eight contained only synonymous substitutions (Fig. 2; Supplementary Table S2).

**Table 2.**
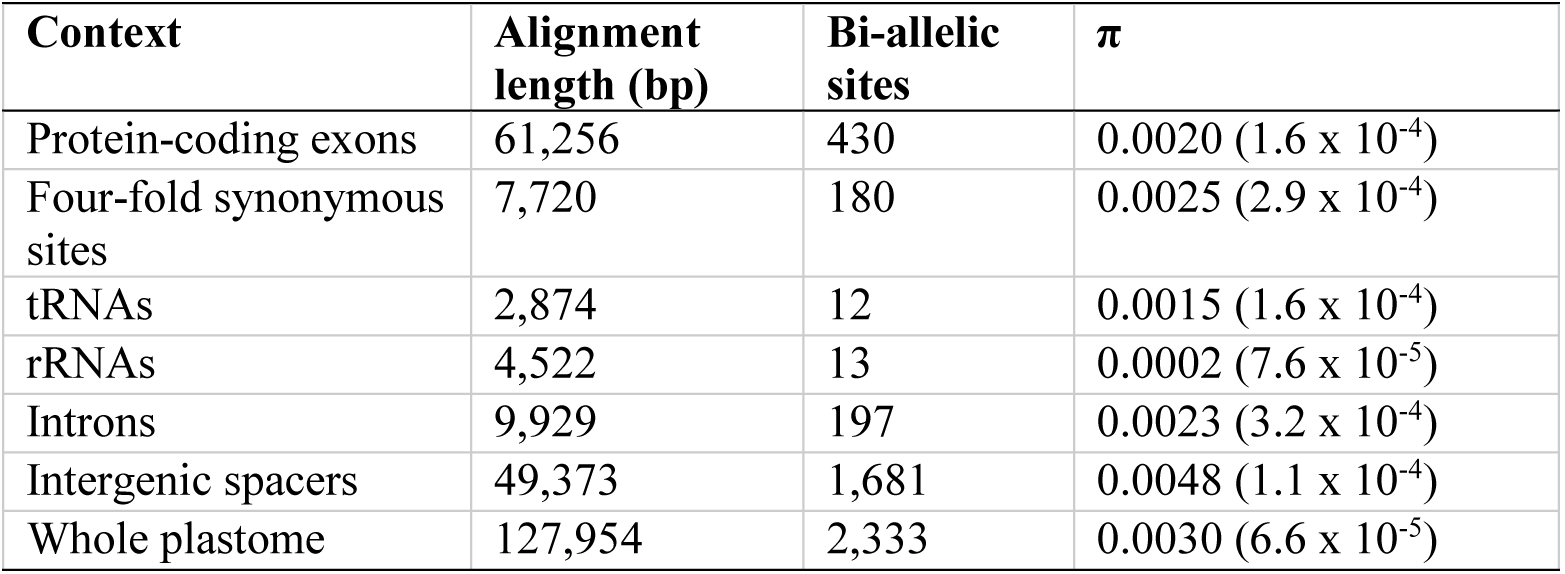
Alignment length, number of polymorphic sites, and nucleotide diversity (π) by genomic context estimated from 65 *Picea* plastomes. Standard errors are reported in parentheses.

| Context | Alignment length (bp) | Bi-allelic sites | $\pi$ |
| --- | --- | --- | --- |
| Protein-coding exons | 61,256 | 430 | 0.0020 ($1.6 \times 10^{-4}$) |
| Four-fold synonymous sites | 7,720 | 180 | 0.0025 ($2.9 \times 10^{-4}$) |
| tRNAs | 2,874 | 12 | 0.0015 ($1.6 \times 10^{-4}$) |
| rRNAs | 4,522 | 13 | 0.0002 ($7.6 \times 10^{-5}$) |
| Introns | 9,929 | 197 | 0.0023 ($3.2 \times 10^{-4}$) |
| Intergenic spacers | 49,373 | 1,681 | 0.0048 ($1.1 \times 10^{-4}$) |
| Whole plastome | 127,954 | 2,333 | 0.0030 ($6.6 \times 10^{-5}$) |

**Figure 2.**
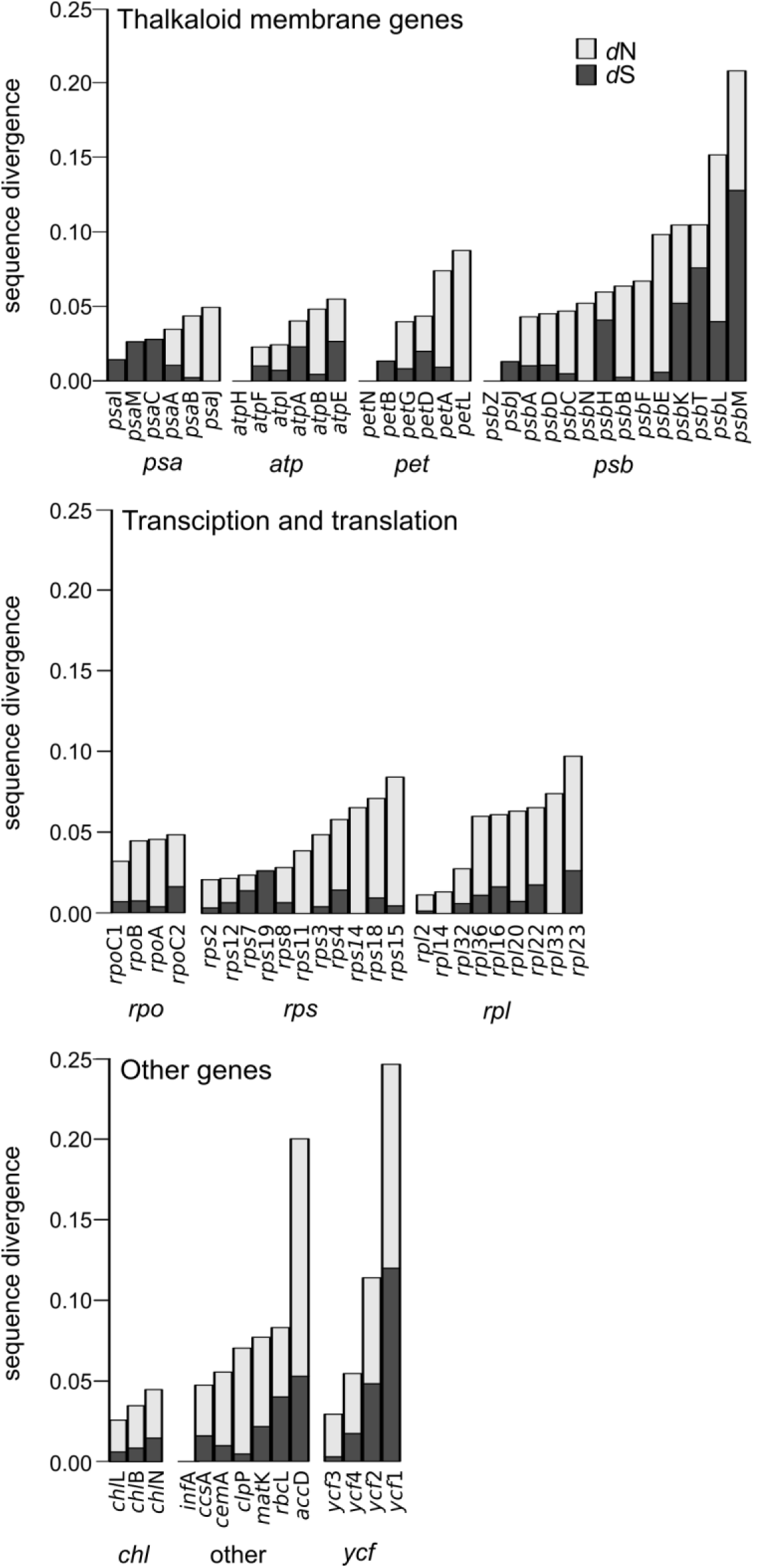
Synonymous (*d*S) and non-synonymous (*d*N) substitution rates for *Picea* plastome genes grouped by complex and function. Rates were estimated using the maximum likelihood method implemented in PAML (Yang 2007). Substitution rates were significantly *(p* < 0.05) elevated in the photosystem II *(psb)* and hypothetical open reading frame *(ycf)* genes. Per-gene estimates are reported in Supplementary Table S2.

The variation in substitution rates among genes prompted us to test for selection acting on them. Codon-based models of nucleotide substitution strongly supported (p ≤ 0.001) positive evolution (ω > 1) in 12 out of 70 genes (Table 3). Five additional genes were significant at 0.001 < *p* ≤ 0.05 but we did not consider them further (Supplementary Table S2). Most genes had one or two positively selected codons, but the massive *ycf1* gene contained 73 sites (3.5% of the predicted protein) potentially under selection. However, many of these sites were located within repetitive sequences and poor alignment may have generated false positives. After excluding sites in tandem repeats, positive selection was still supported at 21 codons (presented in Table 3).

**Table 3.** Positive selection inferred from the M1a and M2a random site models implemented in PAML *(p* ≤0.001). Positively selected sites have ω > 1 with posterior probability ≥ 0.95 according to the Bayes empirical Bayes method. Amino acids variants observed at each putative selected sites are listed in parenthesis. Site number refers to the position in the *Picea* alignment for that protein.

| Complex | Gene | -ln(M1a) | -ln(M2a) | Sites |
| --- | --- | --- | --- | --- |
| ATP synthase | *atpA* | -2210.70 | -2182.54 | 185(P,L), 232(P,S), 253(K,N), 299(S,L), 314(V,L), 427(A,T) |
|  | *atpE* | -617.32 | -578.11 | 119(P,L) |
| Photosystem I | *psaC* | -352.21 | -342.13 | 35(K,R) |
| Photosystem II | *psbA* | -1517.02 | -1481.65 | 155(T,S,A) |
|  | *psbD* | -1538.38 | -1525.49 | 156(P,S) |
|  | *psbH* | -346.92 | -337.02 | 16(S,A), 21(R,T) |
|  | *psbT* | -198.05 | -168.65 | 30(P,L,Q,S) |
| Large ribosomal subunit | *rpl22* | -683.02 | -664.74 | 7(N,Y,K) |
| RNA polymerase | *rpoC2* | -5556.69 | -5540.32 | 326(N,K), 870(R,I) |
| Small ribosomal subunit | *rps19* | -399.85 | -388.39 | 20(K,Q) |
| Hypothetical open reading frame | *ycf1* | -13094.20 | -12574.89 | 68(A,T), 176(V,A), 502(S,Y), 612(L,I), 619(H,Y), 739(R,K), 959(F,L), 1054(A,S), 1083(L,F), 1139(N,K), 1149(S,A), 1223(F,L), F1306(F,L), 1378(L,F), 1383(S,L), 1507(S,L), 1633(L,F), 1708(D,E), 1709(K,Q), 1790(V,T), 1912(R,E) |
|  | *ycf4* | -815.00 | -799.87 | 118(L,P) |

### Phylogenetic discordance in the *Picea* plastome

Phylogenetic analysis of the 65 whole plastomes using maximum likelihood resolved five major clades comprising: i) the North American *P. engelmannii, P. glauca,* and *P. mexicana* (hereinafter ‘glauca’), ii) *P. abies* and predominately northeast Asian taxa (‘abies’), iii) *P. mariana* and four geographically-disparate species (‘mariana’), iv) taxa from the Qinghai-Tibetan Plateau and surrounding regions (‘likiangensis’), and v) a diverse clade containing species representing the entire geographic range of genus (‘chihuahuana’; Fig. 3). Relationships within most clades were strongly supported with the notable exception of the ‘abies’ clade, in which several nodes had less than 50% support. The uncertain topology is likely due to the rapid and recent radiation of this clade during the last two million years (Lockwood et al. 2013, Ran et al. 2015). In contrast, the backbone topology of the genus was poorly resolved, with 52% of trees supporting a ‘glauca’-‘abies’ sister relationship and 39% supporting a most recent common ancestor of the ‘glauca’, ‘abies’, and ‘mariana’ clades.

**Figure 3.**
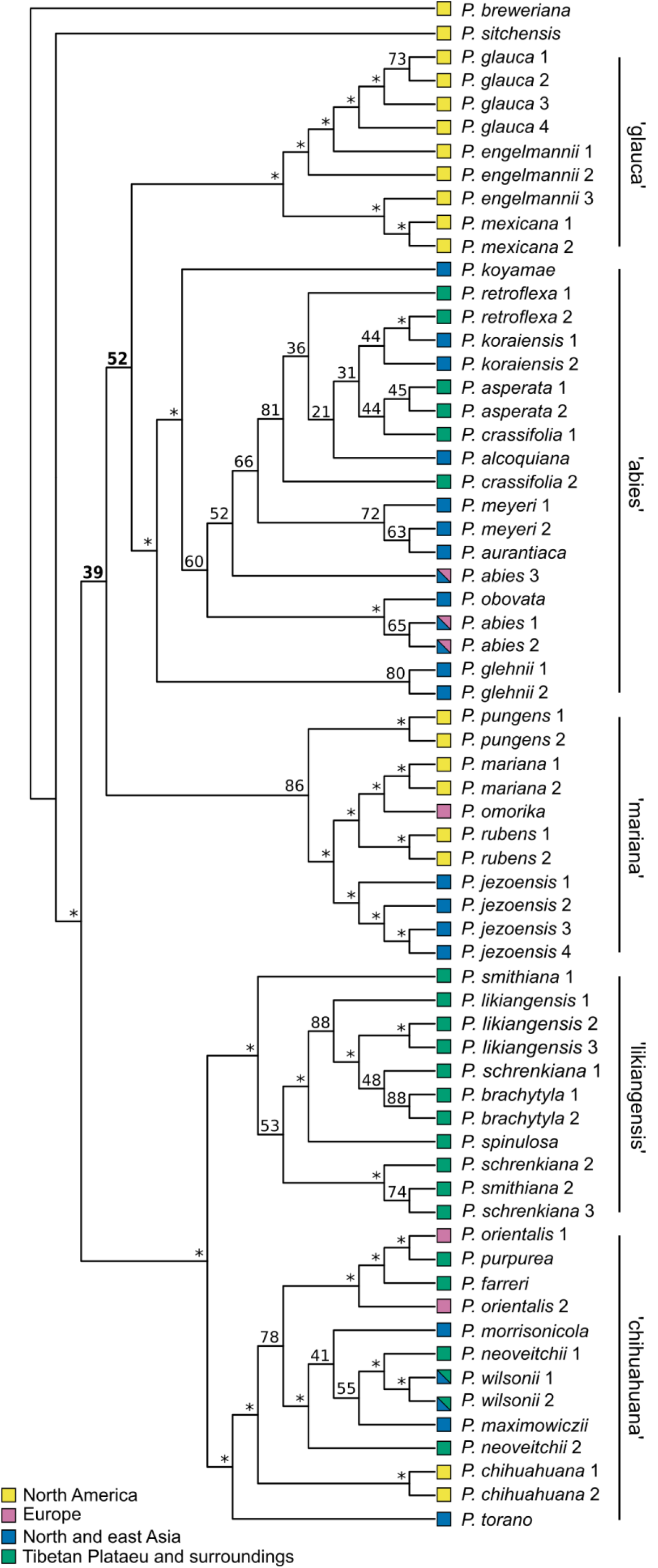
Maximum likelihood analysis of the whole plastome alignment of 65 *Picea* accessions provides strong support for many shallow nodes but the relationships between the ‘glauca’, ‘abies’, and ‘mariana’ clades are unresolved. The majority rules extended consensus tree is presented and node labels denote the proportion of bootstrap replicate trees supporting the bipartition. Nodes with ≥ 90% support are indicated by an asterisk. Colored squares denote the geographic distribution of the species. Numbers after species names correspond to the accession information in Supplementary Table S1.

We analyzed bipartition frequencies in bootstrapped trees to assess the presence of alternate topologies at these two weakly-supported backbone nodes. Out of 1,000 replicates, the ‘glauca’ clade occurred in a bipartition with *P. sitchensis* and *P. breweriana* at even greater frequency than the relationship included on the consensus tree in Fig. 3 (46% vs. 39% of bootstrap replicates). Both the ‘mariana’ and ‘glauca’ clades were grouped with the ‘abies’ clade to the exclusion of all other species with almost equal frequency (52% vs. 48%). This result is consistent with prior plastid-based studies, which discovered conflicting sister clades for ‘abies’ taxa depending on the locus analyzed (cf. Bouillé et al. 2011; Ran et al. 2015). These results suggest the presence of highly-supported yet conflicting topologies in the whole-plastome alignment.

Saturation, selection, or recombination could explain the near-equal occurrence of conflicting bipartitions in the whole-plastome dataset and the disagreement among earlier studies. To assess the causes of phylogenetic discord across the plastome, we estimated phylogenies from five different subsets: 1) protein-coding exons; 2) concatenated intergenic spacers; and 3-5) subsets corresponding to the three major structural regions (F1, F2, and F3) of Pinaceae plastomes, which are subtended by large repeats that likely serve as recombination substrates (Wu et al. 2011; see Fig. 1). We reason these repeats could facilitate recombination between divergent plastome copies if biparental inheritance and plastid fusion occurred. Discordance among structural fragments would be consistent with our hypothesis of recombination, but discordance between genic and intergenic regions would be more consistent with parallel mutations from convergence or saturation.

We implemented randomization tests of Robinson-Foulds (RF) tree distances, quartet distances (QD), and the parsimony-based incongruence length difference (ILD) metric to test the significance of phylogenetic incongruence between *a priori* data subsets. Incongruence between intergenic spacers and protein-coding exons was only supported by the ILD test (Table 4), which potentially has elevated Type I and II error rates under a variety of circumstances (Planet 2006). The lack of strongly-supported discordance between intergenic and coding sites indicates the absence of pervasive differences in phylogenetic signal, even if some sites are genuinely discordant. In contrast, all three tests strongly supported incongruence between the F3 region versus the F1 and F2 regions (Table 4, Fig. 4). While the placement of ‘mariana’ is not highly supported in the F3 region topology, it is excluded from the bipartition comprising the ‘glauca’ and ‘abies’ clades in 99% of bootstrap replicates. In the F1 and F2 phylogenies, ‘mariana’ is resolved as the sister clade of ‘abies’ with high support (Fig. 4; Supplementary Fig. S1). Together, these results suggest different evolutionary histories among the plastome structural regions.

**Table 4.**
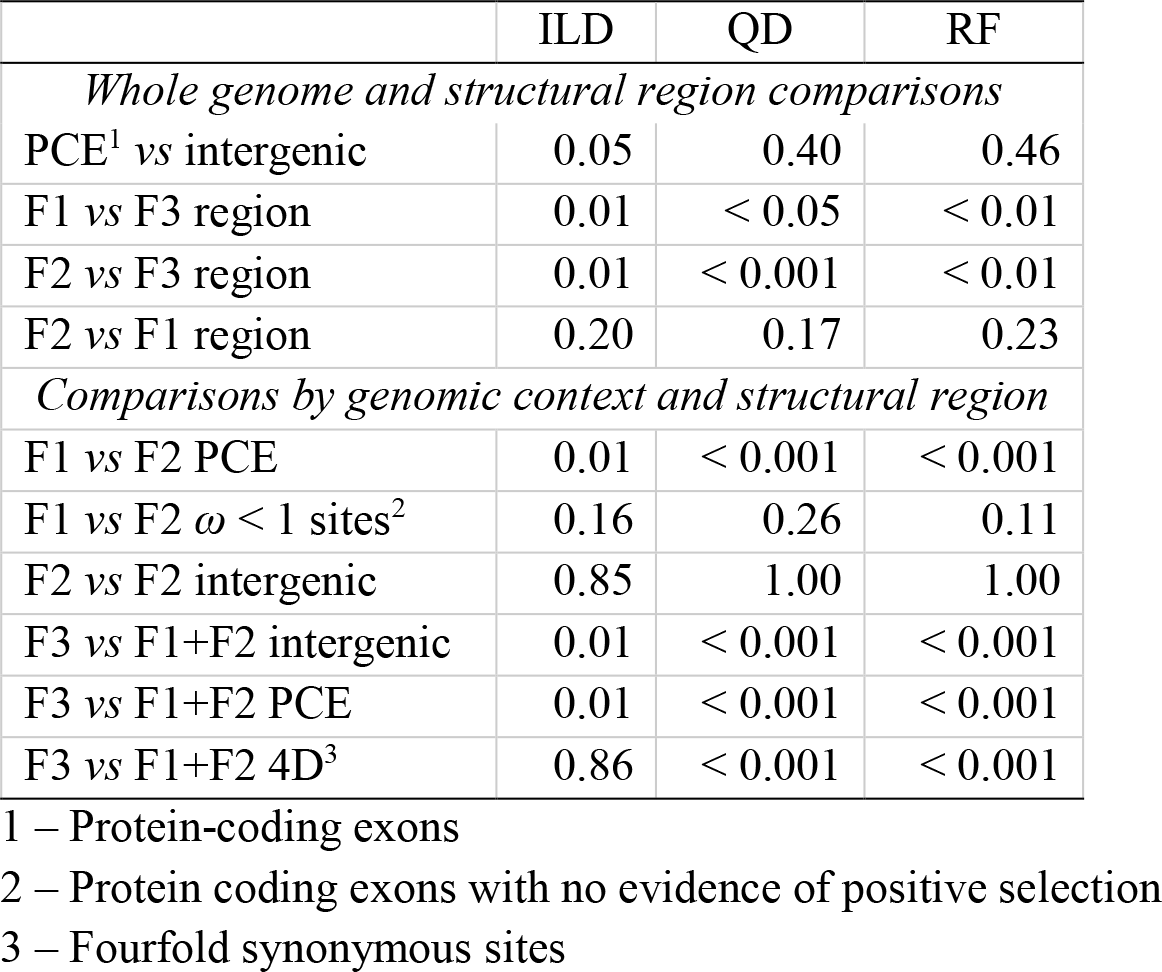
Significance of phylogenetic incongruence between plastome subsets according to the incongruence length difference (ILD), quartet distance (QD), and Robinson-Foulds (RF) metric permutation tests. F1, F2 and F3 regions refer to the three major single copy units of the *Picea* plastome (see Fig. 1). The F1 and F2 regions are concatenated for comparisons by genomic context against the F3 region.

**Figure 4.**
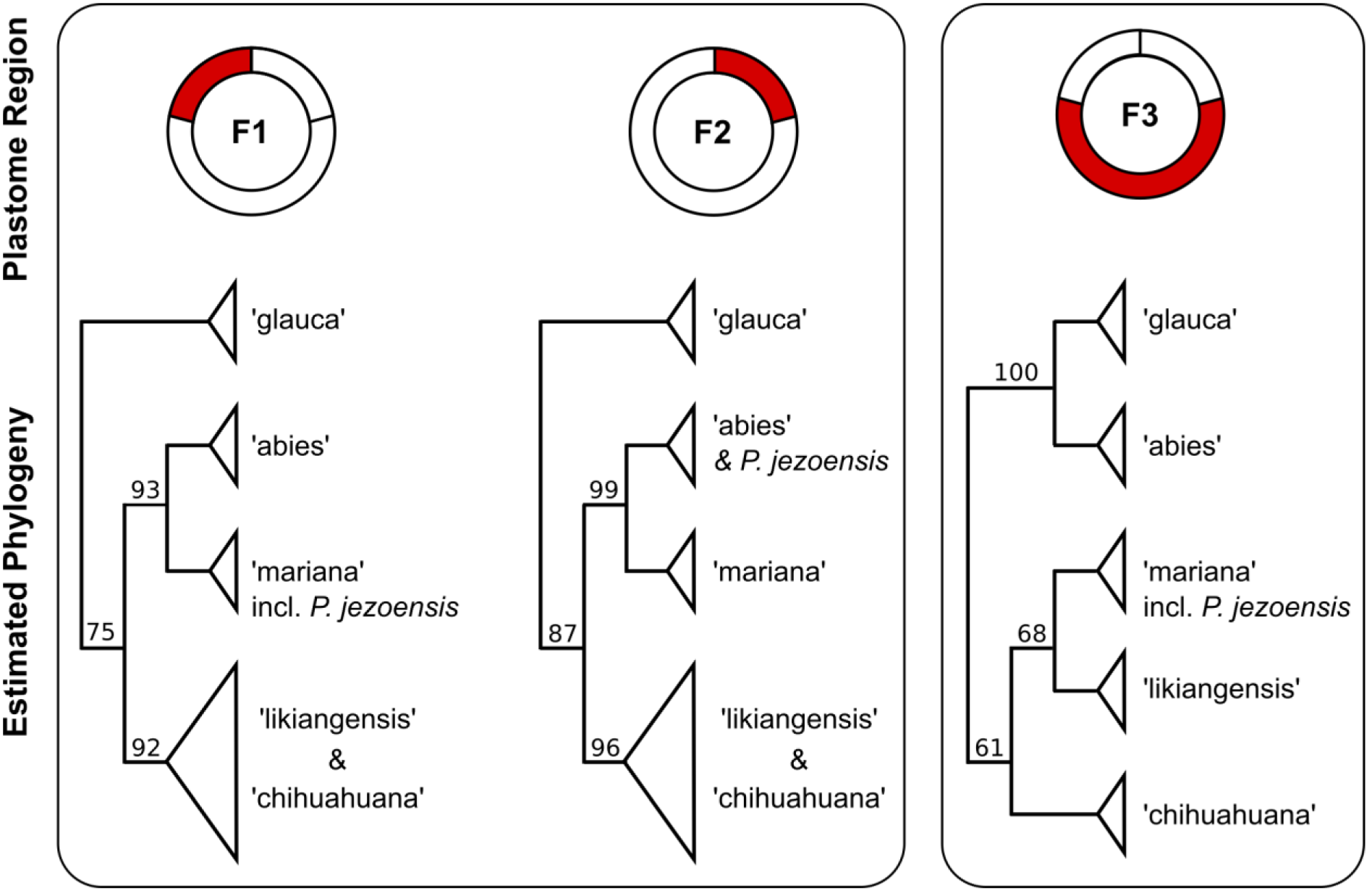
Phylogenies inferred from the structural regions of the *Picea* plastome are strongly discordant according to topology and character based incongruence tests. While the F3 region supports a sister relationship between the ‘glauca’ and ‘abies’ clades, the F2 and F1 regions suggest ‘abies’ and ‘mariana’ share a more recent common ancestor. Full phylogenies are presented in Supplementary Fig. S1.

Differences in gene content, nucleotide diversity, and substitution rates across the three regions could potentially explain the variation in topologies. We tested if phylogenetic discordance between regions could be explained by differences in sequence evolution by comparing only the 1) intergenic sites, 2) protein-coding exons, and 3) four-fold synonymous sites (4D) on each region. Intergenic sites in the F1 and F2 regions yielded statistically indistinguishable topologies but protein-coding exons were significantly discordant (Table 4; Supplementary Fig. S2). Given the paucity of polymorphic 4D sites on these two regions, we instead compared protein-coding exons without significant evidence of selection. A disproportionate number of codons on the F1 region are putatively under positive selection (see Table 3), and their removal yielded topologies no more different than expected by chance (Table 4). As discordance between the F1 and F2 regions can be explained by just eight polymorphisms, we concatenated these two regions for tests against the larger F3 region because using a similar number of sites each phylogeny should improve the accuracy of the permutation tests.

Protein-coding exons, intergenic sites, and 4D sites on the concatenated F1 and F2 regions were strongly discordant with those on the F3 region according to all three tests, with the exception of the ILD test on 4D sites (Table 4). As there are only ~100 parsimony informative 4D sites across the entire plastome, the ILD test may have insufficient power to detect discordance (Planet 2006). Each subset of sites (e.g., 4D sites on the F3 region) yielded a similar topology as the entire region (Fig. 4), albeit with poorer resolution (Supplementary Fig. S2). Therefore, phylogenetic discordance between the F3 versus the F1 and F2 regions encompasses intergenic, synonymous, and non-synonymous sites, and topological differences among the three fragments cannot be adequately explained by saturation or selection.

### Recombination tests

Phylogenetic discordance among structural regions is consistent with our *a priori* hypothesis of interspecific recombination in the plastome. To further support this inference, we employed RDP4 (Martin et al. 2015) to implement unguided tests for recombination on the whole plastome alignment. This two-step approach helps safeguard against potentially spurious results from exploratory tests yet allows the data to refute our hypothesis that recombination should involve the repeats subtending the three structural regions.

The seven tests implemented in RDP4 identified two putative recombination events, both with breakpoints estimated near region junctions (Fig. 5, Table 5). All tests supported a recombinant plastome shared by the ‘abies’ clade originating from parents from the ‘glauca’ and ‘mariana’ clades (Table 5). While the majority of the ‘abies’-type plastome suggested a close relationship with the ‘glauca’ clade, a ~30 kbp segment was highly similar to *P jezoensis,* a species from the ‘mariana’ clade distributed from central Japan to northeastern Siberia. This relationship between the ‘abies’ clade and *P. jezoensis* is also evident in our *a priori* phylogenetic analysis, which supports a shared ancestor of the F2 region (Fig. 4). Maximum likelihood recombination breakpoints were placed near the repeats bounding the F2 region, but broad confidence intervals also encompassed most of the F1 region (Fig. 5). Although the phylogenetic discordance and recombination tests differ somewhat in the extent of the exchange, both support a recombinant plastome in the ‘abies’ clade with at least the F2 region inherited from a ‘mariana’-like parent.

**Figure 5.**
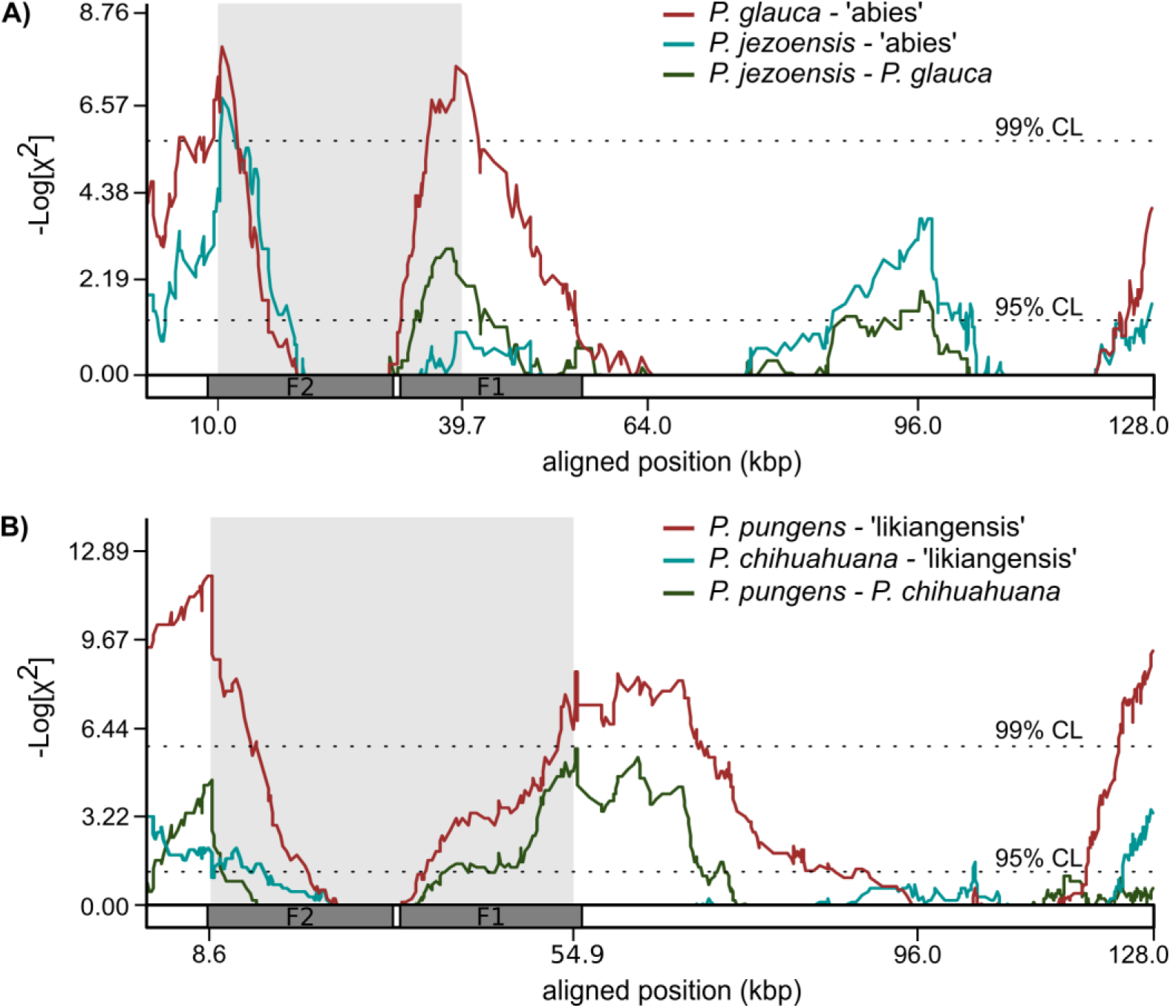
Unguided tests for recombination in the *Picea* whole plastome alignment, as illustrated by the MaxChi method (Maynard Smith 1992). A) All seven recombination tests supported a recombinant plastome shared by the entire ‘abies’ clade originating from interspecific recombination with a *P. jezoensis*--like species. On the y-axis, lines represent the 2x2 contingency χ^2^ value of the number of polymorphic sites on either side of a given window. Breakpoints are inferred to occur where this value maximized and the recombinant region is shaded in grey. A schematic representing the structural components of the plastome is superimposed on the x-axis. B) Five tests supported a recombinant plastome shared by the ‘likiangensis’ clade and estimated breakpoints occurred near the F2 and F1 region junctions.

**Table 5.** Bonferonni corrected p-values for the two recombination events detected in the ‘abies’ and ‘likiangensis’ dades by unguided tests as implemented in RDP4

| Recombination Test | ‘abies’ | ‘likiangensis’ |
| --- | --- | --- |
| RDP | $6.71 \times 10^{-09}$ | $2.36 \times 10^{-5}$ |
| GENECOV | $1.89 \times 10^{-11}$ | n.s. |
| BootScan | $6.71 \times 10^{-5}$ | n.s. |
| MaxChi | $1.81 \times 10^{-7}$ | $6.11 \times 10^{-6}$ |
| Chimaera | $1.83 \times 10^{-4}$ | $6.87 \times 10^{-8}$ |
| SiScan | $9.46 \times 10^{-11}$ | $1.55 \times 10^{-3}$ |
| 3Seq | $4.67 \times 10^{-4}$ | $2.57 \times 10^{-9}$ |

n.s. – not significant

A second putative recombination event spanning the F1 and F2 fragments involved the ‘likiangensis’ clade of Qinghai-Tibetan Plateau taxa and an unknown species similar to *P. pungens* a species from the ‘mariana’ clade (Table 5, Fig. 5). The F3 region topology supported a sister relationship between the ‘likiangensis’ and ‘mariana’ clades with 68% bootstrap support (Fig. 4), whereas the F1 and F2 regions intercalated taxa from the ‘likiangensis’ and ‘chihuahuana’ clades (Fig. 4, Supplementary Fig. 2). As the RF, QD, and ILD randomization tests are global measures of discordance, this more equivocal recombination event likely contributed to the highly significant results in Table 4.

## Discussion

Uniparental inheritance, strong functional constraints, and absent recombination are key but often untested assumptions in plant evolutionary studies (Wolfe & Randle 2004). Using whole genome assemblies and comprehensive taxon sampling, we found that *Picea* plastomes are highly conserved in sequence and structure but shaped by interspecific recombination and punctuated by putative positive selection. Our results add to the growing body of evidence of complex plastid evolution and demonstrate how overly simple models can lead to misleading inferences.

### Sequence divergence

Plastome substitution rates are typically low in land plants (Wolfe et al. 1987). Consequently, interspecific distances in the *Picea* plastome are of the same magnitude as intraspecific variation in Pinaceae nuclear genes (e.g. Mosca et al. 2012). Genome-wide substitution rates are not commonly reported in genus-level whole plastome studies, however, substitution rates in *Picea* are similar to those reported from more limited sets of loci in *Pinus* (Willyard et al. 2007; Wang & Wang 2014).

Both genera have ancient origins in the Cretaceous (Wang et al. 2000; Klymiuk & Stockey 2012), but the diversification of extant pine and spruce species occurred within the last 20 million years, perhaps facilitated by a cooling climate after the Miocene optimum (Willyard et al. 2007; Lockwood et al. 2013; Wang & Wang 2014; Ran et al. 2015). Although we did not estimate absolute substitution rates explicitly, the similarity of the *Pinus* and *Picea* relative substitution rates and divergence times together suggests a similar per-generation mutation rate in these two Pinaceae genera.

### Selection on *Picea* plastid genes

Positive selection on plastid genes has been widely inferred in angiosperms (e.g. Carbonell-Caballero et al. 2015; Hu et al. 2015) and in some cases potentially adaptive changes have been demonstrated at the protein or phenotype level (Mengistu et al. 2000). Gymnosperm plastomes are less studied. An analysis of *Pinus* plastomes found more limited evidence for positive selection (6/71 genes; Parks et al. 2009), in contrast to the 20% of *Picea* genes apparently affected. Only two genes, *psbH* and *ycf1,* were inferred to have codons with ω > 1 in both *Pinus* and *Picea.* To our knowledge, psbH has not been widely implicated in positive evolution in angiosperms, and high ω at this gene may represent a unique attribute of Pinaceae. In contrast, *ycf1* is exceptionally divergent across land plants (Dong et al. 2015) and shows elevated ω in some angiosperm lineages (e.g. Hu et al. 2015). Surprisingly, *ycf1* encodes an essential component of plastid protein import apparatus (Kikuchi et al. 2013), the hypervariable repeat motifs of which may serve an adaptive function (de Vries et al. 2015).

Consistent with an adaptive explanation, positively selected sites in *Picea* were disproportionately inferred in genes involved in photosynthesis (Table 3). However, non-synonymous mutations may become fixed through non-adaptive processes. Small effective population sizes, rare recombination, and strong mutational bias are all characteristic of plastomes and can facilitate the fixation of slightly deleterious alleles (Kimura 1983; Felsenstein 1974; Moran 1996). On the other hand, plastids have highly efficient DNA repair systems, which could be expected to purge deleterious mutations (Day & Madesis 2007; Maréchal & Brisson 2010). Site-directed mutagenesis and in vitro characterization of mutant proteins (e.g. Lan et al. 2013) could help clarify the role of adaptive and non-adaptive processes in shaping plastid substitution rates.

### Reticulate evolution and chimeric plastomes

Recombination between divergent haplotypes has been invoked to explain phylogenetic discordance among plastid loci (Erixon & Oxelman 2008; Bouillé et al. 2011; D’Alelio & Ruggiero 2015). Although recombination has been demonstrated in somatic hybrids (e.g. Medgyesy et al. 1985), evidence from natural populations is rare and based on limited data (Marshall et al. 2001; Huang et al. 2001; Erixon & Oxelman 2008; Bouillé et al. 2011; D’Alelio & Ruggiero 2015). Using hypothesis and data-driven analyses, we found evidence for at least one interspecific recombination event in *Picea* plastomes. Although a robust coalescent species tree is not yet available for *Picea,* topologies inferred for the F3 plastome region closely match those from mitochondrial loci (Ran et al. 2015) and nuclear loci (Zou et al. 2016; J-Q Liu, unpublished data), which suggests the ‘glauca’ and ‘abies’ clades may be related by vertical descent. In contrast, the F2 and potentially the F1 region of the ‘abies’ plastome was inherited horizontally from a member of the ‘mariana’ clade, most likely *P. jezoensis* or an extinct relative. Formation of chimeric plastomes via interspecific recombination would require at a minimum: 1) hybridization, 2) temporary breakdown of uniparental inheritance, and 3) plastid fusion and intermolecular recombination.

Like many plant taxa, extant *Picea* are widely interfertile and many ‘abies’ species successfully hybridize with the ‘mariana’ clade in controlled crosses (Wright 1955). In natural populations, many parapatric spruces hybridize readily (e.g. Jaramillo-Correa et al. 2005; Du et al. 2011), including distantly-related species (Hamilton & Aitken 2013). Comparison of phylogenies inferred from mitochondrial and plastid loci, which are differentially inherited in conifers, also support widespread introgression throughout the genus (Bouillé et al. 2011; Ran et al. 2015). In particular, *P. jezoensis* has a ‘mariana’-type plastome but more closely resembles the ‘abies’ clade at mitochondrial and some nuclear loci (Ran et al. 2015). Fossil and genetic evidence indicate the continuous presence of *P. jezoensis* and its extinct relative *P. hondoensis* in northeastern Russia throughout the Pleistocene (Aizawa et al. 2007). The most recent common ancestor of the ‘abies’ clade likely emerged in eastern Asia during this same time period (Ran et al. 2015), and introgression between a *P. jezoensis-like* species and the ‘abies’ ancestor may have occurred here, prior to the clade’s diversification and colonization of Eurasia.

Mechanisms maintaining uniparental inheritance are varied (Bock 2007), but hybridization or interpopulation crosses have been linked to a breakdown of these processes (Hansen et al. 2007; Ellis et al. 2008; Barnard-Kubow et al. 2017). In *Helianthus* and *Passiflora,* rates of plastid leakage vary by family and species but realized biparental inheritance occurs in ~2-4% of offspring overall, respectively (Hansen et al. 2007; Ellis et al. 2008). As biparental inheritance is not uncommon on an evolutionary timescale, fusion of the two parental plastids is probably the limiting step preventing more widespread recombination (Day & Madesis 2007). To our knowledge, plastid fusion has not been observed directly in natural populations but has been demonstrated experimentally in somatic hybrids (e.g. Medgyesy et al. 1985). Once fused, recombination could occur between highly similar genomes through normal DNA replication and repair mechanisms (Day & Madesis 2007; Maréchal & Brisson 2010). Spread of the chimeric plastome through the ancestor of the ‘abies’ clade may have then proceeded through entirely neutral processes, such as allele surfing during range expansion (Excoffier & Ray 2008).

The prevalence of sexual recombination in plastomes is unknown. Taxa with higher rates of biparental plastid inheritance, including groups prone to hybridization (Hansen et al. 2007; Ellis et al. 2008; Barnard-Kubow et al. 2017), are presumably the most susceptible. In addition, three genera with predominately paternal plastid inheritance, including *Picea,* have been suggested to form chimeric plastomes (Marshall et al. 2001; Huang et al. 2001), which may indicate a link between this inheritance mode and a propensity for recombination. However, until further research clarifies the factors promoting sexual plastome recombination, adding recombination tests to phylo- and population genetic pipelines is prudent.

### A word of caution for ‘chloroplast phylogenomics ‘

Plastome phylogenies effectively estimate the history of a single gene and do not necessarily represent species histories. Incomplete lineage sorting can cause the plastome to vary markedly from the species tree, and introgression can further exacerbate this discordance (Doyle 1992). While whole *Picea* plastomes yielded high resolution within most clades, they also revealed conflicting histories generated by vertical descent and introgression. Furthermore, the three genomes in *Picea* suggest incongruent histories (Bouillé et al. 2011; Ran et al. 2015), and substantial gene-tree discordance is also evidence among nuclear loci (Ran et al. 2015). Reconstructing a useful approximation of the species tree for *Picea* – and likely many other plant genera - will likely necessitate methods explicitly allowing for gene tree heterogeneity arising from both incomplete lineage sorting and introgression.

Plastome haplotypes for many species were not reciprocally monophyletic, an issue which further confounds the use of single-gene phylogenies as species tree proxies. Haplotype sharing can arise from some combination of incomplete lineage sorting, hybridization, cryptic and poorly-delimited species, or misidentified material. For example, Norway spruce (*P. abies)* was polyphyletic in our dataset, with the two Scandinavian plastomes more closely related to *P. obovata* than to a Norway spruce originating from Serbia. A polyphyletic Norway spruce was noted by Lockwood et al. (2013) in their predominately mitochondrial dataset, which indicates a need to clarify species boundaries in this taxon. In the absence of a robust species tree estimate, we urge caution when interpreting plastome phylogenies and note results from exemplar accessions may not generalize to an entire species.

## Conclusions

We used whole plastome sequences and comprehensive taxon sampling to characterize sequence divergence, positive selection, and evolutionary histories in *Picea.* Most significantly, we found chimeric plastomes formed by ancient introgression and recombination between divergent clades. Chimeric plastomes explain the conflicting results of earlier plastid-based studies in *Picea*: phylogenies inferred from different loci alternately reflect vertical descent and reticulate evolution. As whole plastome sequences become increasingly available, we suggest future studies utilize structural features, such as repeats, to develop phylogenetic hypotheses to test for recombination. Given that hybridization (Ellstrand et al. 1996; Whitney et al. 2010) and the potential for biparental plastid are common in plants (Corriveau & Coleman 1988; Zhang et al. 2003), interspecific plastome recombination may not be unique to *Picea*.

## Materials and Methods

### Plant material

A total of 61 representatives from the 35 commonly recognized *Picea* species (Farjon 1991; Eckenwalder 2009) were sourced from natural populations and cultivated collections. Multiple accessions were sequenced for broadly distributed species or those with spatially-disjunct or fragmented ranges. For cultivated trees, we preferred individuals obtained as seed or seedlings from natural populations and selected wild accessions outside of known hybrid zones whenever possible (see Supplementary Table S1 for provenances and voucher numbers). Additional complete chloroplast genomes for *P. abies* (Nystedt et al. 2013), *P. glauca* (Jackman et al. 2015), *P. morrisonicola* (Wu et al. 2011), and *P. jezoensis* (Yang et al. 2016) were downloaded from Genbank (Supplementary Table S1).

### Genome sequencing and assembly

Total genomic DNA was isolated from leaf material using the Macherey-Nagel NucleoSpin^®^ Plant II kit according to the manufacturer’s instructions (Düren, Germany). Paired-end libraries with ~350 bp inserts were constructed for 77 accessions for this and another study and sequenced on Illumina HiSeq 2000 and 2500 lanes using 2×100 bp and 2×125 bp chemistry, respectively, at the National Genomics Infrastructure hosted at Science for Life Laboratory (SciLifeLab, Stockholm, Sweden). Paired-end libraries with ~350 bp inserts for 17 accessions for this and another study were multiplexed on a single Illumina HiSeq 4000 lane at the University of Texas at Austin (Austin, Texas, USA) using 2×150 bp chemistry. Finally, an entire lane of 2×100 bp reads from the Illumina MiSeq platform was available for one accession. We developed a *de novo* assembly strategy by first comparing assemblies produced by SPAdes v. 3.7.0 (Bankevich et al. 2012; Prjibelski et al. 2014) using either total genomic reads or reads pre-filtered for similarity to the plastome. For read filtering, we used local alignment in bowtie2 v. 2.2.7 (Langmead & Salzberg 2012) using the *P. abies* plastome as a reference and tested both the ‘very fast’ and ‘very sensitive’ presets. The most contiguous assemblies with the highest depth of coverage were obtained by using pre-filtered reads and results were nearly identical for both local alignment presets (results not shown). Therefore, we used ‘very fast’ local alignment followed by assembly using the ‘careful’ parameter, default error correction and kmer selection, and a minimum contig size of 500 bp. Sequencing methods, assembly metrics, and Genbank accession numbers for each accession reported in Supplementary Table S1.

### Annotation and alignment

We tested for structural rearrangements in *de novo* scaffolds of each accession versus the *P. abies* reference genome using nucmer, which were visualized as dot plots with mummerplot as contained within the software package MUMmer v. 3.0 (Kurtz et al. 2004). The scaffolds of each accession were aligned against the *P. abies* reference and pseudomolecules were generated in Geneious v. 9.0.4 for further analysis. Multiple sequence alignment of pseudomolecules was conducted in MAFFT v. 7.245 (Katoh & Standley 2013) using the fast Fourier transformation approximation option, a partition size of 1,000 and three iterative refinements. Sequences were annotated according to the *P. abies* reference genome using the BLAST-like ‘transfer annotations’ tool in Geneious using an identity cut-off of 60%. All annotations were manually curated to ensure accurate identification of genes, pseudogenes, and reading frames. Plastome maps were created with CIRCOS (Krzywinski et al. 2009).

### Sequence polymorphism and divergence

Nucleotide diversity (π) was estimated using the R package PopGenome v. 2.1.6 (Pfeifer et al. 2014) by genomic context and in 350 bp sliding windows. Standard errors were estimated using 500 bootstrap replicates. Average synonymous (*d*S) and non-synonymous (*d*N) substitution rates were estimated per-gene using the maximum likelihood method (Goldman & Yang 1994) implemented in the codeml program in PAML v. 4.9 (Yang 2007). We inferred natural selection at the codon-level from the ratio of *d*N to *dS* (ω) using nested random site models (M1a & M2a; Wong et al. 2004; Yang et al. 2005) as implemented in codeml. The maximum-likelihood tree inferred from concatenated protein-coding exons was used as the input tree with relationships between the five major clades and any branches with < 70% bootstrap support constrained to polytomies. Both copies of *ycf12* and *psbI* were excluded because they are located within large repeats (white arrows in Fig. 1) and were not uniformly recovered across all accessions.

### Phylogenetic analyses

Sequence alignments were prepared for phylogenetic analysis by stripping sites with more than 20% missing data and removing tandem repeats using Phobos v. 3.3.12 ( <http://www.ruhr-uni-bochum.de/spezzoo/cm/cmphobos.htm>). We used PartitionFinder v. 1.1.1 to identify best-fit partitioning schemes of nucleotide evolution (Lanfear et al. 2012, 2014). We defined an initial model giving each protein-coding gene and codon position a partition, and tRNAs, rRNAs, introns were each assigned to a single partition, respectively. We constrained the model of nucleotide evolution to GTR+Γ and used the linked branch lengths model. Partition searches were performed using the ‘rcluster’ clustering algorithm with the default percentage. Optimal partitioning schemes identified using the whole genome data were subsequently applied to each of the five data subsets: 1) protein-coding exons, 2) concatenated intergenic spaces, and 3-5) the three structural regions (F1, F2, and F3) of the *Picea* plastome. Phylogenies were inferred from the partitioned datasets in RAxML v. 8.2.4 (Stamatakis 2014) using rapid bootstrapping under the autoMRE convergence criterion and were summarized on an extended majority rule consensus tree (Aberer et al. 2010). Trees are rooted by *P. breweriana* for display, which has been inferred to be the sister lineage to all other extant spruces in previous studies (Bouillé et al. 2011; Lockwood et al. 2013; Ran et al. 2015).

We tested for phylogenetic discordance using character and tree-based methods. The character-based ILD test was performed using PAUP* v. 4.0a149 (Swofford 2002) using 100 replicates with a heuristic starting seed, random-stepwise additions, and nearest-neighbor interchange branch swapping with the search limited to 100,000 re-arrangements for computational feasibility. Permutation tests based on tree QD (Bryant et al. 2000) and RF (Robinson & Foulds 1981) distances tested if phylogenies estimated from different *a priori* subsets are more different than expected by chance. To simulate tree distances under the null hypothesis, we calculated QD and RF values between 10,000 trees estimated from randomly sampled plastome sites. Sets of random sites were selected to be comparable in size to the subsets of interest. Then, we calculated RF and QD distances between 10,000 randomly selected pairs of trees sampled from the bootstrap replicates of the *a priori* subsets. We implemented the permutation test using the R package ‘perm’ (Fay & Shaw 2010) using 2,000 replications and estimated *p*-values using the default algorithm. QDs were calculated using the program rqtdist v. 1.0 (Sand et al. 2013) and RF distances in the R packages ‘phangorn’ (Schliep 2011) and ‘ape’ (Paradis et al. 2004).

### Recombination tests

Unguided tests for recombination were conducted in the RDP4 package (Martin et al. 2015) using the RDP (Martin & Rybicki 2000), GENECOV, BootScan (Salminen et al. 1995), MaxChi (Maynard Smith 1992), Chimaera (Posada & Crandall 2001), SiScan (Padidam et al. 1999), and 3seq (Boni et al. 2007) methods. After identifying a recombination event, RDP4 implements a hidden Markov model to estimate breakpoint positions. Then, RDP4 identifies the probable recombinant sequence using the PHYLPRO (Weiller 1998), VISRD (Lemey et al. 2009), and EEEP (Beiko & Hamilton 2006) methods. Bonferrroni corrections were applied to set the family-wise error rate to 0.05. We tested the whole-plastome alignment for recombination and an alignment stripped of gaps and poorly aligned regions but the results did not differ. In addition, altering window sizes and indel processing options did not alter the results so default values were used for the final runs.

## Acknowledgements

We are grateful to Daniel Luscombe (Bedgebury National Pinetum), Peter Brownless (Royal Botanic Garden Edinburgh) Charles Keith (Charles Keith Arboretum), Erik Dahl Kjær and Albin Lobo (University of Copenhagen, Denmark), Piotr Banaszczak (Rogów Arboretum), Johnny Schimmel (Arboretum Norr), Dan Crowley (Westonbirt, The National Arboretum), László Csiba and Rhinaixa Duque-Thüs (Royal Botanic Gardens, Kew), Jaakko Saarinen (Mustila Arboretum), and Jianquan Liu (Lanzhou University and Sichuan University) for contributing samples to this study. We acknowledge support from Science for Life Laboratory and the National Genomics Infrastructure, Sweden for providing assistance in massive parallel sequencing. Computations were performed on resources provided by SNIC through Uppsala Multidisciplinary Center for Advanced Computational Science (UPPMAX) under Project SNIC 2015-6-67. Douglas Scofield at UPPMAX and Nicolas Delhomme at the Umeå Plant Science Center (UPSC) Bioinformatics Facility provided bioinformatics support. This study was supported by the Trees and Crops for the Future (TC4F) program. ARS was supported in part by Bergvik Skog AB. Constructive criticism from anonymous reviewers improved the manuscript.

## Supplementary material

Supplementary Table S1. Locality and accession information for all 65 included *Picea* plastomes. Assembly statistics are reported for the 61 *de novo* assemblies.

Supplementary Table S2. Per gene estimates of nucleotide diversity (π), *d*N, *d*S, and codon-based maximum likelihood tests for positive selection.

Supplementary Figure S1. Extended majority rules consensus trees estimated from the entire sequence of the F1, F2 and F3 regions, respectively. Nodes with 100% bootstrap support are denoted with an asterisk.

Supplementary Figure S2. Extended majority rules consensus trees estimated from fourfold synonymous sites, intergenic spaces, and protein coding exons from each region (F1, F2, and F3), respectively. Nodes with 100% bootstrap support are denoted with an asterisk.

